# Genomic signatures accompanying the dietary shift to phytophagy in polyphagan beetles

**DOI:** 10.1101/399808

**Authors:** Mathieu Seppey, Panagiotis Ioannidis, Brent C. Emerson, Camille Pitteloud, Marc Robinson-Rechavi, Julien Roux, Hermes E. Escalona, Duane D. McKenna, Bernhard Misof, Seunggwan Shin, Xin Zhou, Robert M. Waterhouse, Nadir Alvarez

## Abstract

**Background:** The diversity and evolutionary success of beetles (Coleoptera) are proposed to be related to the diversity of plants on which they feed. Indeed the largest beetle suborder, Polyphaga, mostly includes plant-eaters among its ~315,000 species. In particular, plants defend themselves with a diversity of specialized toxic chemicals. These may impose selective pressures that drive genomic diversification and speciation in phytophagous beetles. However, evidence of changes in beetle gene repertoires driven by such interactions remains largely anecdotal and without explicit hypothesis testing.

**Results:** To address this, we explored the genomic consequences of beetle-plant trophic interactions by performing comparative gene family analyses across 18 species representing the two most species-rich beetle suborders. We contrasted the gene contents of species from the mostly plant-eating suborder Polyphaga with those of the mainly predatory Adephaga. We found gene repertoire evolution to be more dynamic, with significantly more adaptive lineage-specific expansions, in the more speciose Polyphaga. Testing the specific hypothesis of adaptation to plant-feeding, we identified families of enzymes putatively involved in beetle-plant interactions that underwent adaptive expansions in Polyphaga. There was especially strong support for the selection hypothesis on large gene families for glutathione S-transferase and carboxylesterase detoxification enzymes.

**Conclusions:** Our explicit modeling of the evolution of gene repertoires across 18 species identifies adaptive lineage-specific gene family expansions that accompany the dietary shift towards plants in beetles. These genomic signatures support the popular hypothesis of a key role for interactions with plant chemical defenses, and for plant-feeding in general, in driving beetle diversification.

## Introduction

Species richness among eukaryotes varies substantially, with some clades having only a few representatives and others comprising hundreds of thousands of extant species. In particular, the class Insecta outnumbers all other classes with more than half of all described extant species (Farrell, 1998; Grimaldi and Engel, 2005). Beetles (Coleoptera) encompass approximately 380,000 described species, representing ca. 40% of described insect diversity (Slipinski et al., 2011). Several hypotheses have been proposed to explain this richness, notably their complex interactions with flowering plants (Farrell, 1998; Leschen and Buckley, 2007; McKenna et al., 2009, 2015; Zhang et al., 2018) and a high lineage survival rate (Hunt et al., 2007). Nevertheless, detailed supporting evidence from molecular genetic studies remains sparse, making it difficult to assess the relative importance of these and other potentially important contributing factors (Barraclough et al., 1998; Suchan and Alvarez, 2015).

The remarkable evolutionary success of beetles may have been driven by the interplay between their trophic niche and their genomic content and architecture. This is based on the premise that environmental and ecological conditions are likely to be predominant factors influencing the fate of genetic variation in populations under natural selection (Barrick and Lenski, 2013), eventually driving divergence into distinct species (Seehausen et al., 2014). Among all components of the biotic environment, the trophic niche (principal source of nourishment) of an organism plays a crucial role in shaping the evolution of phenotypic innovations and their underlying genomic changes, e.g. feeding modes in cichlid fishes (Parsons et al., 2016), mouth development in *Pristionchus* nematodes (Ragsdale et al., 2013), and bitter taste receptors in vertebrates (Li and Zhang, 2014). Among several hypotheses explaining the tremendous diversity among beetles, a shift from an ancestral diet as saprophages (detritus-feeding) or mycophages (fungi-feeding) (Betz et al., 2003) to phytophagy (feeding on living plant material in a broad sense) is often evoked (Farrell, 1998; Leschen and Buckley, 2007; McKenna et al., 2009). While the suborder Adephaga (~45,000 species) comprises mostly predatory species, including ground beetles and diving beetles, the largest beetle suborder, Polyphaga (~315,000), is predominantly comprised of phytophagous clades, among which the most species-rich families are weevils (Curculionidae, ~51,000), longhorn beetles (Cerambycidae, ~30,000), and leaf beetles (Chrysomelidae, ~32,000) (Slipinski et al., 2011). Phytophagy appeared approximately 425 million years ago, quickly after terrestrial life was established (Labandeira, 2002). It progressively diversified to target most plant tissues (Labandeira, 2013), shortly before the radiation of flowering plants 120-100 million years ago (Grimaldi, 1999). In response, plants have evolved diverse strategies to protect themselves, which in turn impose selective pressures on the animals that feed on them.

While many biological processes are likely to play a role in this evolutionary battle, a key weapon in the arsenal of phytophagous insects’ adaptations is their ability to neutralize or minimize the effects of plant secondary compounds. Protein families known to be crucial for eliminating harmful plant toxins are cytochrome P450 monooxygenases (P450s), carboxylesterases (CEs), UDP-glycosyltransferases (UGTs), and glutathione S-transferases (GSTs) (Voelckel and Jander, 2014). While P450s and CEs modify residues to make compounds more hydrophilic, UGTs and GSTs conjugate xenobiotic compounds to hydrophilic molecules. Detoxification is completed by membrane transporters, such as ATP-binding cassette (ABCs) transporters, which move xenobiotic compounds to where they can either be excreted, or less frequently sequestered in order to be reused as a defense mechanism (Voelckel and Jander, 2014). Additionally, to prevent phytophagous insects from digesting their tissues, plants produce enzyme inhibitors that block catalytic sites or compete with the substrates of enzymes involved in digestion. The major families affected are endopeptidases, such as cysteine (CYSs), and serine (SERs) proteases, as well as more specific enzymes such as glycoside hydrolases (GHs), certain types of which are able to break down polysaccharide molecules, including cellulose, hemicellulose and pectin in plant cell walls (McKenna et al., 2016; Pauchet et al., 2010). Other adaptations to phytophagy include repertoires of chemoreceptors that are crucial for finding appropriate food sources (Goldman-Huertas et al., 2015), and the specialization of mouthparts in response to plant mechanical barriers, which are highly diversified in insects (Labandeira, 1997).

As lineages diverge, their genomes accumulate changes, some of which are expected to be directly linked to functional adaptations. Identifying such genomic features and linking them to phenotypic differences, whilst robustly distinguishing between the effects of stochastic changes and natural selection (Hurst, 2009), is critical to deciphering the genomic drivers of species radiations (Shaw and Lesnick, 2009). Changes include point substitutions, which may affect existing functional elements, but also larger-scale changes such as duplications, from individual genes to entire genomes, which by adding new members to the repertoires of key gene families may constitute an ideal mechanism to facilitate the emergence of novel functions leading to successful phytophagy (Kondrashov, 2012). Whereas newly generated gene copies are usually redundant or deleterious and pseudogenized, rendering the gene copy non-functional (Innan and Kondrashov, 2010), they are sometimes maintained. Particularly interesting cases of gene family expansions are the ones restricted to specific lineages, resulting in lineage specific expansion (LSE). Evolutionary mechanisms causing LSE are numerous and not all adaptive (see Innan and Kondrashov, 2010 for a comprehensive review). However, duplicated gene copies may provide an immediate selective advantage and be maintained by selection. This can be due to an increased dosage of the gene, or to changes following the duplication being selected in one gene copy but not the other, which might allow evolution towards a different function in so-called neo-functionalization processes. Enzymes are considered particularly relevant candidates for such evolutionary processes as they could expand their range of substrates (Francino, 2005).

Here we apply a comparative genomics approach to examine the evolution of genes putatively involved in plant-insect interactions by sampling from the two largest beetle suborders, which, generally-speaking, present contrasting trophic niches. We contrast exemplars from the characteristically predaceous Adephaga with exemplars from Polyphaga and we hypothesize that plant-insect interactions during the dietary shift to phytophagy should be accompanied by genomic evolutionary signatures visible at the subordinal scale. Using genomic and transcriptomic data from 18 beetle species, we estimate ancestral gene family content, taking into account gene gains and losses across the species phylogeny, to identify significant LSEs of gene families related to phytophagy and signatures of adaptive expansions in these families. Ignoring sensory receptors, as their evolution might be driven by agents other than those related strictly to trophic niche (Brito et al., 2016), and morphological genes, as their inferred association with diet is less robust, we focus on genes coding for enzymes, for which adaptive LSE specific to Polyphaga would suggest a role for detoxification and digestive pathways in driving adaptation and speciation.

## Results

### A representative sampling of the two major coleopteran suborders

Reliable estimation of gene gain and loss events requires a robust evolutionary framework, i.e. a phylogeny that includes the species studied, as well as the characterization of gene families across complete gene sets from these same species. To study adaptation to phytophagy, we sampled from both Adephaga (mostly predaceous) and Polyphaga (with diverse trophic habits, including a very large number of phytophagous species). A balanced sampling of each suborder was achieved comprising twelve transcriptomes and six genomes, with Benchmarking Universal Single-Copy Ortholog (BUSCO) completeness estimates (Simão et al., 2015; Waterhouse et al., 2018) ranging from 71.9% to 97% (Figure 1, Table 1). The species phylogeny was estimated using 405 BUSCO genes found to be complete in all species and the strepsipteran outgroup, *Stylops melittae* (Figure 1). Protein-coding sequence predictions ranged from 9,844 to 24,671 genes per beetle species. These sequences matched 14,908 Arthropoda orthologous groups (OGs) containing at least one species of Coleoptera in the OrthoDB v8 catalog (Kriventseva et al., 2015). This represented a minimum of 6,742 and a maximum of 11,149 OGs for *Carabus frigidus* and *Leptinotarsa decemlineata*, respectively. OGs containing genes from only one of the two sampled suborders were excluded, resulting in a total of 9,720 OGs for the analysis that have evolutionary histories traceable to the last common ancestor of beetles. Functional annotations of the sequences within these OGs were used to identify and assign several of them to enzyme families relevant to the tested hypothesis. These candidate OGs comprised four UGTs, 22 P450s, 19 CEs, six GSTs, four SERs, seven CYSs, 28 ABCs, and one GH, for a total of 91 candidate OGs from eight families of genes (i.e. functional categories) (Table 2).

**Figure 1.**
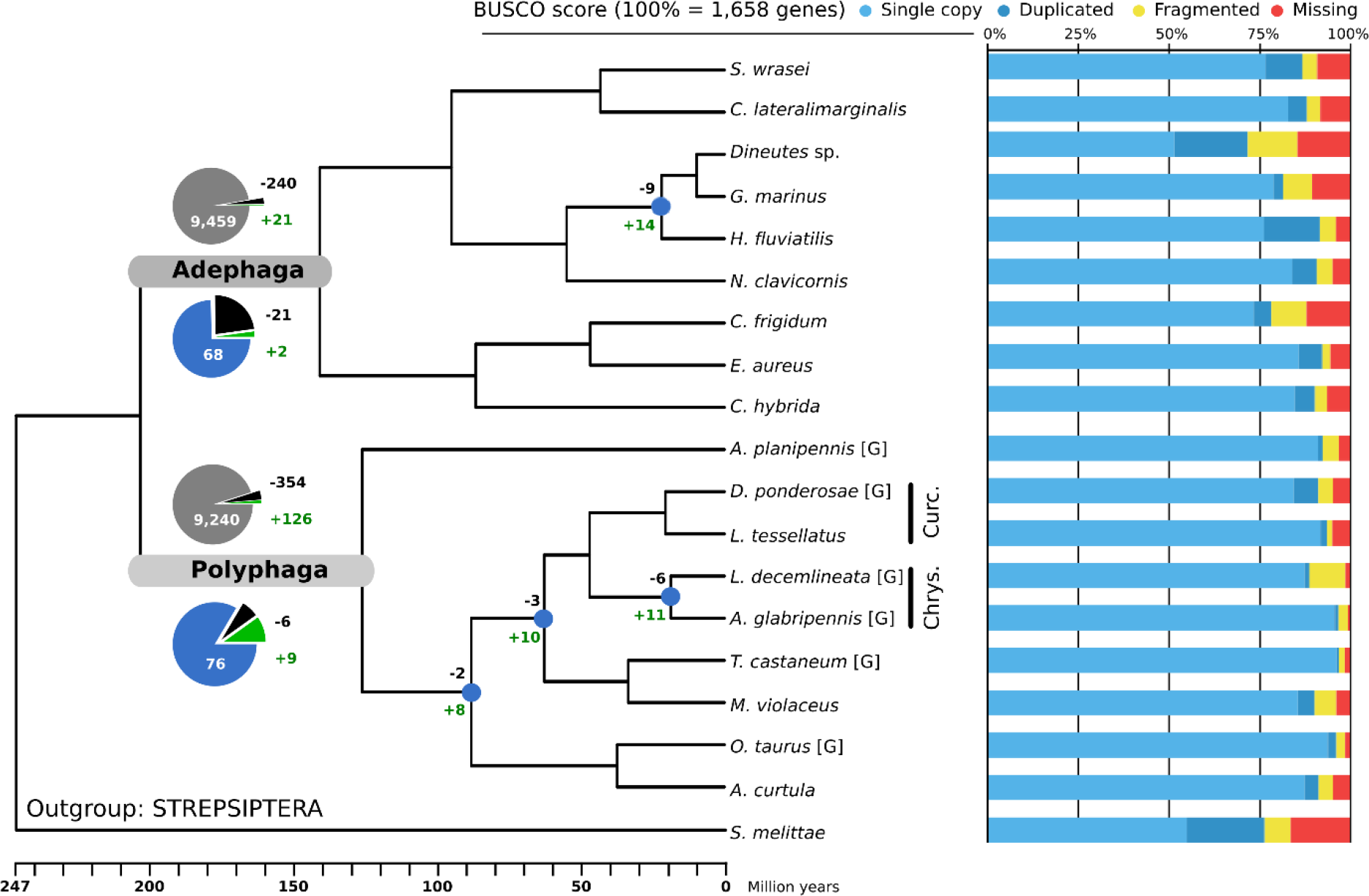
The ultrametric species phylogeny with gene family expansions and contractions quantified for nodes of interest and bar charts showing completeness of the genomic and transcriptomic datasets studied. The species tree was built from 405 single-copy orthologs and constrained to have Geadephaga (*C. frigidum*, *E. aureus*, *C. hybridia*) and Hydradephaga (the six other Adephaga) as monophyletic sister clades (e.g., following McKenna et al., 2015). Branch lengths are scaled in millions of years. Maximum likelihood bootstrap support was 99 or 100% for all branches. [G] symbol indicates data from species with sequenced genomes with the remaining species being from transcriptomes. The numbers of orthologous groups (OGs) with expansions (+) and contractions (−) are displayed at the root node of each suborder. Pie charts show proportions of OGs with gene losses (black) and gene gains (green) with respect to OGs with no significant losses or gains for all considered OGs (grey) and only the candidate OGs (blue). While gains constitute only a small subset of all OGs in both suborders, the proportion of gains is much larger among candidate OGs in Polyphaga. The nodes indicated by blue circles in the Polyphaga sub-tree lead to species-rich clades containing species that are largely phytophagous (e.g., Chrysomelidae and Curculionidae, respectively Chrys. and Curc.) and experienced larger proportions of gains among the candidate OGs. The Benchmarking Universal Single-Copy Ortholog (BUSCO) scores indicate the relative levels of completeness and putative gene duplications for the genome-based and transcriptome-based datasets in terms of 1,658 BUSCOs from the insecta_odb9 assessment dataset.

**Table 1.** Beetle genomes and transcriptomes included in the study. Taxonomic classifications are listed with data sources, as well as completeness (Benchmarking Universal Single-Copy Ortholog, BUSCO, score, C=Complete, S=Complete Single-Copy, D=Complete Duplicated, F=Fragmented, M=Missing), number of predicted proteins, and number of orthologous groups (OGs) with genes from each species. The outgroup species used in the phylogeny, *Stylops melittae*, belongs to the order Strepsiptera, which is the sister group of Coleoptera (Niehuis et al., 2012).

| Species | Short form | Suborder | Family | Type of assembly | Accession | Bioproject | Source | BUSCO score (1658 genes) | Predicted proteins | OrthoDB groups (OG) |
| --- | --- | --- | --- | --- | --- | --- | --- | --- | --- | --- |
| *Cicindela hybrida* | CHYBR | Adephaga | Carabidae | Transcriptome | GDMH01000000 | PRJNA286505 | 1KITE, this study | C:90.4[S:84.9%,D:5.5%],F:3.4%,M:6.2% | 13,916 | 8,111 |
| *Calosoma frigidum* | CFRIG | Adephaga | Carabidae | Transcriptome | GDLF01000000 | PRJNA286499 | 1KITE, this study | C:78.3[S:73.5%,D:4.8%],F:9.8%,M:11.9% | 9,844 | 6,742 |
| *Elaphrus aureus* | EAURE | Adephaga | Carabidae | Transcriptome | GDPI01000000 | PRJNA286520 | 1KITE, this study | C:92.4[S:85.9%,D:6.5%],F:2.2%,M:5.4% | 12,808 | 8,020 |
| *Noterus clavicornis* | NCLAV | Adephaga | Noteridae | Transcriptome | GDNA01000000 | PRJNA286561 | 1KITE, this study | C:90.9[S:84.1%,D:6.8%],F:4.4%,M:4.7% | 12,981 | 7,918 |
| *Haliplus fluvialis* | HFLUV | Adephaga | Halplidae | Transcriptome | GDMW01000000 | PRJNA286525 | 1KITE, this study | C:91.8[S:76.3%,D:15.5%],F:4.3%,M:3.9% | 19,408 | 8,528 |
| *Cybister lateralmarginalis* | CLATE | Adephaga | Dytiscidae | Transcriptome | GD LH01000000 | PRJNA286512 | 1KITE, this study | C:88.2[S:83.1%,D:5.1%],F:3.7%,M:8.1% | 13,916 | 7,256 |
| *Sinaspidytes wrasei* | SWRAS | Adephaga | Aspidytidae | Transcriptome | GDNH01000000 | PRJNA286492 | 1KITE, this study | C:87.0[S:76.7%,D:10.3%],F:4.0%,M:9.0% | 13,392 | 7,721 |
| *Dineutes sp.* | DINEU | Adephaga | Gyrinidae | Transcriptome | GDNB01000000 | PRJNA286516 | 1KITE, this study | C:71.9[S:51.6%,D:20.3%],F:13.6%,M:14.5% | 14,644 | 7,089 |
| *Gyrinus marinus* | GMARI | Adephaga | Gyrinidae | Transcriptome | GAUY01000000 | PRJNA219564 | 1KITE, Misof et al., 2014 | C:81.5[S:79.0%,D:2.5%],F:8.1%,M:10.4% | 13,867 | 7,663 |
| *Aleochara curtula* | ACURT | Polyphaga | Staphylinidae | Transcriptome | GATW01000000 | PRJNA219522 | 1KITE, Misof et al., 2014 | C:91.4[S:87.5%,D:3.9%],F:4.0%,M:4.6% | 20,280 | 8,513 |
| *Anoplophora glabripennis* | AGLAB | Polyphaga | Cerambycidae | Genome | GCF_000390285 | PRJNA167479 | I5k, McKenna et al., 2016 | C:96.9[S:95.8%,D:1.1%],F:2.7%,M:0.4% | 22,035 | 10,959 |
| *Agrilus planipennis* | APLAN | Polyphaga | Buprestidae | Genome | GCF_000699045 | PRJNA230921 | I5k, unpublished | C:92.5[S:91.2%,D:1.3%],F:4.5%,M:3.0% | 15,497 | 9,089 |
| *Dendroctonus ponderosae* | DPOND | Polyphaga | Curculionidae | Genome | GCF_000355655 | PRJNA162621 | Keeling et al., 2013 | C:91.2[S:86.0%,D:5.2%],F:4.1%,M:4.7% | 13,457 | 8,518 |
| *Leptinotarsa decemlineata* | LDECE | Polyphaga | Chrysomelidae | Genome | GCF_000500325 | PRJNA171749 | I5k, Schoville et al., 2018 | C:88.9[S:87.5%,D:1.4%],F:9.9%,M:1.2% | 24,671 | 11,149 |
| *Laparocerus tessellatus* | LTESS | Polyphaga | Curculionidae | Transcriptome | 10.5281/zenodo.1336288 | N/A | This study | C:93.8[S:91.9%,D:1.9%],F:1.4%,M:4.8% | 18,448 | 8,616 |
| *Meloe violaceus* | MVIOL | Polyphaga | Meloidae | Transcriptome | GATA01000000 | PRJNA219578 | 1KITE, Misof et al., 2014 | C:90.3[S:85.6%,D:4.7%],F:5.9%,M:3.8% | 14,295 | 8,480 |
| *Onthophagus taurus* | OTAUR | Polyphaga | Scarabaeidae | Genome | GCF_000648695 | PRJNA167478 | I5k, unpublished | C:96.2[S:93.9%,D:2.3%],F:2.5%,M:1.3% | 17,483 | 9,315 |
| *Tribolium castaneum* | TCAST | Polyphaga | Tenebrionidae | Genome | GCF_000002335 | PRJNA12540 | Tribolium Genome Sequencing Consortium et al., 2008 | C:97.0[S:96.5%,D:0.5%],F:1.6%,M:1.4% | 16,645 | 9,429 |
| *Stylops melittae* | SMELI | Outgroup | Stylopidae | Transcriptome | GAZM02000000 | PRJNA219603 | 1KITE, Misof et al., 2014 | C:76.5[S:55.0%,D:21.5%],F:7.1%,M:16.4% | 13,026 | 6,104 |

**Table 2.** Candidate gene categories with the keywords and identifiers used to select them from the full sets of sequences annotated with InterProScan. To be included as candidate orthologous groups (OGs) in the category, OGs were required to have at least one sequence matching both a UniRef and an InterProScan entry, and an additional gene ontology term in the case of serine proteases.

| Gene Family Category | InterProScan (Pfam or InterPro identifiers) or Gene Ontology | UnifRef KeyWord | Number of OGs |
| --- | --- | --- | --- |
| UDP-glycosyltransferases (UGTs) | PF00201 | name:"cluster UDP glucuronosyltransferase"<br>OR name:"cluster UDP glycosyltransferase" | 4 |
| Cytochrome P450 oxidases (P450s) | PF00067 | name:"cluster Cytochrome P450" | 22 |
| Carboxylesterases (CEs) | PF02230, PF00135 | name:"cluster carboxylesterase"<br>OR name:"carboxylic ester hydrolase" | 19 |
| Glutathione S-transferases (GSTs) | PF00043, PF02798 | name:"cluster Glutathione S-transferase" | 6 |
| Serine proteases (SERs) | PF00450, PF12146, PF05577, GO:0008236 | name:"cluster Serine protease"<br>OR name:"cluster Serine peptidase" | 4 |
| Cysteine proteases (CYSs) | PF00112 | name:"cluster cysteine protease"<br>OR name:"cluster cystein protease"<br>OR name:"cluster Papain" | 7 |
| ABC transporters (ABCs) | PPF00005, PF00664 | name:"cluster ABC" | 28 |
| Glycoside hydrolases (GHs) | IPR000334, IPR000743, IPR001360, IPR001547 | name:"cluster Glycoside hydrolase" | 1 |
| **Total** |  |  | 91 |

### Polyphaga exhibit more frequent gains across a larger set of OGs

Analysis of per-species gene counts of the complete set of 9,720 OGs was performed with the Computational Analysis of gene Family Evolution (CAFE v3) (Han et al., 2013) tool. The mode considering distinct gene gain (λ=0.0019 gain/gene/million years) and gene loss (μ=0.0018 loss/gene/million years) was preferred over a single value for λ and μ, having a significantly greater maximum likelihood score (see Methods). The λ (gain) and μ (loss) values predicted when CAFE was run on each suborder separately were λ=0.0020 and μ=0.0027 for Adephaga, versus λ=0.0023 and μ=0.0021 for Polyphaga, showing a tendency for Adephaga to lose genes and for Polyphaga to gain genes. Among the 9,720 OGs were 21 with reported expansions originating at the Adephaga root and 126 at the Polyphaga root (see Figure 1 to locate the nodes). Conversely, 240 OGs showed gene losses for Adephaga and 354 for Polyphaga. Two expansions and 21 losses affected the candidate OGs for Adephaga, and nine expansions and six losses for Polyphaga. Other polyphagan nodes leading to phytophagous-rich clades (i.e., Chrysomeloidea and Curculionidae) also exhibited more candidate OGs expanding than contracting (Figure 1). All counts of gene gains and losses per node are presented in Supplementary Figures 1 and 2. Additionally, CAFE assigned individual OG p-values of < 0.01 to a subset of 910 (9.3%) OGs, which, according to De Bie et al., 2006, indicates gene families likely to have experienced accelerated rates of gain and loss. These are interesting to investigate further as they may represent large OGs of potentially unequal size between the suborders. Among these were 26 of the 91 candidate OGs (28.6%), a significantly larger proportion (2-sample test for equality of proportions, chi-square test, p-value < 0.0001) compared with just 9.3% of non-candidate OGs.

### Signatures of adaptive expansion are more prevalent in Polyphaga

All 910 OGs with significant variations in their gene content were tested for signatures of adaptive expansion in each suborder, by comparing Brownian motion (BM, neutral) to Ornstein-Uhlenbeck (OU, selective pressure) evolutionary models (Beaulieu et al., 2012). As mentioned previously, these included 26 OGs that belong to one of the functional categories listed in Table 2 (“candidate” OGs). The models consider per-species gene count as a trait that can evolve towards a value, which may or may not differ between the two suborders and may or may not be guided by selective pressure; we call this the “optimum” value in models integrating selection. In total, 21 OGs displayed a higher optimum for Adephaga (0.2% of the initial 9,720 OGs) and 88 for Polyphaga (0.9%). Eight of these 88 polyphagan OGs (Table 3 and gene trees in Figure 2 and Supplementary Figures 3-9) are candidate OGs belonging to one of the candidate gene families of Table 2, while none of the 21 adephagan OGs belong to any of the candidate gene families. The proportion of OGs with expansions and higher optima in the background (all “candidate” and remaining “control” OGs) was significantly larger for Polyphaga compared to Adephaga (2-sample test for equality of proportions, chi-squared, 88/9720 vs. 21/9720, p-value < 1e-09), indicating that Polyphaga have experienced globally more LSE under selection on protein-coding genes. Furthermore, a test for enrichment (see Methods) of OGs with LSE under selection from the candidate families (Table 2) compared to the background was significant for Polyphaga (8/91 vs. 88/9720, p-value < 1e-09). The same test applied individually on each candidate gene family within the candidate dataset demonstrated that categories enriched for LSE under selection in Polyphaga were GSTs (3/6 positive tests, fdr-corrected p-value < 1e-09) and CEs (3/19, fdr-corrected p-value < 0.005), as shown in detail in Table 4.

**Figure 2.**
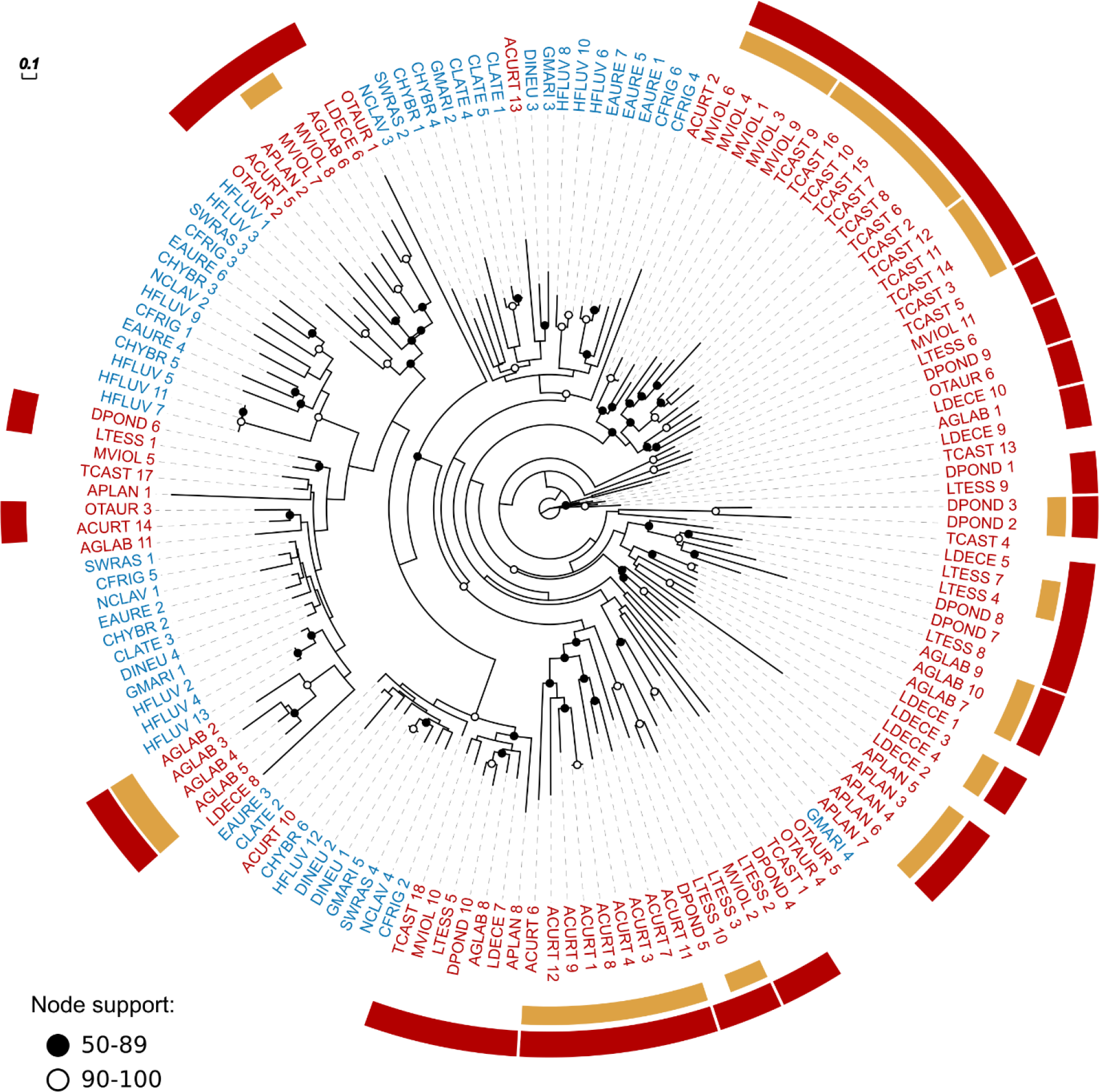
Molecular phylogeny from the largest glutathione S-transferase (GST) orthologous group among those exhibiting lineage-specific expansions driven by selection. Red labels indicate genes belonging to species of Polyphaga, accounting for 98 out of 152 genes (their Ornstein-Uhlenbeck per-species optimum is 11.69 vs. 6.85 for Adephaga (blue labels), see Table 3). The presence of several clades of polyphagan and adephagan genes delineates duplication events following the divergence of the two suborders. Encircling the gene labels are red bars that highlight polyphagan clades with bootstrap support of >50% and yellow bars that highlight intra-specific duplications with bootstrap support of >50%. Corresponding full names of species are given in Table 1. Branch lengths represent substitutions per site and bootstrap support below 50% is not displayed.

**Table 3.** Candidate orthologous groups (OGs) with CAFE overall p-values p<0.01 for which a model favoring selection for larger sizes in Polyphaga showed a greater likelihood. OG identifiers for functional category cytochrome P450s (P450), carboxylesterases (CE), glutathione S-transferases (GST), and cysteine proteases (CYS) are from OrthoDB v8 (ODB8 ID). Small-sample-size corrected Akaike Information Criterion (AICc) values are reported for all tested models. BM1 (Brownian motion with a single rate for the whole tree), BMS (Brownian motion with different rates for each regime), OU1 (selection towards the same optimum for both regimes), all representing the null hypothesis. OUM (selection towards two optima, same variance) and OUMV (selection towards two optima, two variances), representing the alternative hypotheses. The mean values in each suborder (Adephaga versus Polyphaga) are presented in the last two columns. Values in bold font indicate the preferred (maximum likelihood) model. A delta AICc > 2 is required for H1 to be retained.

| Category | ODB8 ID | BM1<br>AICc<br>H0.1 | BMS<br>AICc<br>H0.2 | OU1<br>AICc<br>H0.3 | OUM<br>AICc<br>H1.1 | OUMV<br>AICc<br>H1.2 | Mean<br>Adephaga | Mean<br>Polyphaga |
| --- | --- | --- | --- | --- | --- | --- | --- | --- |
| P450 | EOG805VG7 | 148.37 | 153.21 | 143.35 | 148.10 | **137.95** | 34.13 | 34.47 |
| CE | EOG87DCWX | 143.23 | 143.77 | 143.63 | **138.91** | 141.15 | 6.55 | 18.78 |
| CE | EOG8KD911 | 87.08 | 91.23 | 82.90 | 89.10 | **79.74** | 0.89 | 2.86 |
| CE | EOG876NDC | 80.64 | 85.67 | 80.08 | 87.42 | **77.23** | 1.72 | 3.48 |
| GST | EOG87WR3Z | 86.24 | 87.76 | 76.05 | 74.40 | **72.12** | 1.71 | 3.16 |
| GST | EOG81RS7Z | 108.77 | 114.44 | 109.19 | 113.76 | **103.79** | 6.85 | 11.69 |
| GST | EOG85F05D | 117.62 | 115.88 | 111.53 | 107.62 | **106.32** | 5.69 | 9.16 |
| CYS | EOG8JDKNM | 91.85 | 91.62 | 89.74 | **85.66** | 88.25 | 1.80 | 3.78 |

**Table 4.** Gene family category and candidate orthologous group (OG) enrichments among positive results. The top panel presents the statistical significance of each test for enrichment of candidate gene families among the positive results when compared to the background, for Polyphaga. The lower panel indicates the number of positive results in both suborders, for candidate OGs and background. Significant values at the 0.05 threshold are shown in bold.

| Category | Positive/Total OGs | Category enrichment FDR |
| --- | --- | --- |
| P450 | 1/22 | 0.36268 |
| CE | 3/19 | **0.00252** |
| GST | 3/6 | **0.00016** |
| CYS | 1/7 | 0.16627 |
| UGT | 0/4 | 1.00000 |
| SER | 0/4 | 1.00000 |
| ABC | 0/28 | 1.00000 |
| GH | 0/1 | 1.00000 |
| Category | Positive/Total OGs | Candidate vs. background enrichment p-value |
| Background (Polyphaga) | 88/9,720 | **0** |
| Candidates (Polyphaga) | 8/91 |  |
| Background (Adephaga) | 21/9,720 | 1 |
| Candidates (Adephaga) | 0/91 |  |

## Discussion

Comparative genomic analyses often highlight expanded gene families and link these expansions to biological functions peculiar to, or of special interest in, their focal organism(s). However, these analyses usually do not explicitly test for any hypothesized evolutionary model that might support such links. Here we test a specific hypothesis of adaptation to a phytophagous diet, by comparing candidate gene family repertoires from nine adephagan (a mostly predaceous suborder) and nine polyphagan (a highly phytophagous suborder) beetle species. These candidate families are putatively involved in detoxification of plant allelochemicals and digestion of plant tissues. Specifically, we identify evidence for potentially adaptive gene family expansions in the species rich Polyphaga. This result is robust to potentially confounding factors that could arise from combining genomic and transcriptomic datasets, conservative definitions of candidate gene families, or the greater species richness of the Polyphaga (see discussion points below and Supplementary Information). Through explicitly testing for adaptive LSEs, these results offer conclusive supporting evidence for the key evolutionary role of the phytophagous trophic niche in driving gene family expansions in Coleoptera (specifically Polyphaga), a feature that likely facilitated adaptation of polyphagan beetles to specialized plant feeding.

### Dataset heterogeneity

For the comparison of gene repertoires between the two groups to be unbiased, the gene content of all analyzed species should be of similar accuracy and completeness. The number of predicted proteins for the genomic resources for each beetle species (Table 1, mean 15,977 and standard deviation 3,748) was within the range expected of insects (see Waterhouse, 2015). The average total gene count for Adephaga species (all transcriptomes) was about 4,200 fewer than for Polyphaga, which include two genomes with more than 22,000 genes. This difference in average gene counts is reduced to just 1,384 when considering only genes assigned to the 9,720 OGs selected for the analysis. Our conservative orthology filtering therefore ensured that the comparisons focused on gene families with reliably traceable evolutionary histories that span both groups of beetles. Secondly, assessments of completeness showed that the majority of the datasets contained more than 90% of complete BUSCOs (Figure 1, Table 1). While the dynamically evolving families that are the focus of this study are clearly not universal single-copy orthologs, the high levels of BUSCO completeness support the assumption that the datasets represent good coverage of the species’ gene content. Re-analyses of our data that exclude the two adephagan beetle species with fewer than 80% complete BUSCOs reduced the power of the model tests but nevertheless still identified the three GST OGs that favor a model with a higher optima for Polyphaga (see Supplementary Results). Three of the adephagan transcriptomes showed more than 10% of duplicated BUSCOs, which could have arisen from suboptimal filtering of the transcriptomes, i.e. failure to remove alternative transcripts of the same gene. While such potentially inflated gene counts for these adephagans might prevent the identification of some true expansions in Polyphaga, they do not invalidate those that were identified. Finally, half of the OGs representing positive results showed a higher mean value for polyphagan transcriptomes than genomes, including the three GST OGs (Supplementary Table 1), and explicitly testing for effects due to using both genome and transcriptome data for the species of Polyphaga, by performing a modified OUwie analysis with data type as the regime under selection, identified only one CE (EOG8KD911) for which the favored model linked gene family expansion to species with genomes (see Supplementary Results).

### Candidate OG identification

The annotation strategy was designed to link OGs to candidate gene families based on manually selected keywords used to filter sequence search results, as well as Pfam and InterPro identifiers (Table 2), with the aim of excluding false positives (see Methods). This conservative strategy may not have fully captured all possible candidate OGs, which would therefore have remained in the background set of OGs that were used as controls. For example, we identified nine GST OGs (six were retained as candidates after filtering) while ten subclasses have been identified in arthropods (Roncalli et al., 2015). While the strict (conservative) strategy we employed to identify candidate OGs may have resulted in an underestimate of the extent of the observed effects, this does not invalidate those that were identified. In addition, filtering the OGs to retain only those with genes from both Adephaga and Polyphaga excluded from the analyses any genes that were specific to either suborder. These might include genes with key roles in phytophagy, e.g. enzymes acquired by horizontal gene transfer identified from the *A. planipennis*, *A. glabripennis*, and *D. ponderosae* genomes (McKenna et al., 2016). While acknowledging their importance, here we explicitly tested for adaptive LSE in one lineage versus the other so gene evolutionary histories were required to span the two suborders and thus be traceable to their last common ancestor.

### The more speciose Polyphaga exhibit more dynamic gene repertoire evolution

The μ and λ values reported by CAFE on all 9,720 OGs are consistent with assessments of other insect clades (Hahn et al., 2007; Neafsey et al., 2015). Although the overall gain rate is slightly higher than the loss rate, the number of OGs losing genes reported by CAFE at each individual node is generally larger than the number of OGs with gains. This is reconciled by considering that across Coleoptera many OGs lost a few genes while few families gained many genes. As most OGs display a low number of genes per species, i.e. they are evolving under ‘single-copy control’ (Waterhouse et al., 2011), losing more than one ortholog per species is understandably rare, while there is no theoretical limit for an OG to gain new members. Comparing the two clades, Polyphaga has a higher rate of gene gain and six times more OGs with gains, and while the Adephaga rate of gene loss is higher, Polyphaga have 1.5 times more OGs that have experienced gene losses. Hence, the gene repertoires of Polyphaga exhibit a more dynamic evolutionary history with more gains (rate) in more OGs (counts) and fewer losses (rate) spread out over more OGs (counts). It is possible that this greater dynamism may be generally linked to the greater species richness of Polyphaga, with no specific role for phytophagy underpinning this trend. However, among the candidate OGs for detoxification and digestion there are also more gains in Polyphaga, and, in contrast to the background, fewer losses. Thus, both gain and maintenance are higher for candidates in Polyphaga, which is consistent with a key role for phytophagy in driving dynamic gene repertoire evolution, and particularly LSEs.

### Evidence for adaptive expansions of gene families involved in detoxification in polyphagan beetles

In addition to observing more expansions among candidate OGs in the suborder Polyphaga, the positive results from the OUwie analysis support the hypothesis that selective pressures drive detoxification enzymes towards larger gene family sizes. This is especially pronounced for GSTs, for which half of the OGs tested positive and for which a significant enrichment compared to the background was found. The importance of GSTs in dietary shifts to phytophagy has been noted in mustard-feeding flies, where duplicated GSTs involved in the mercapturic acid pathway showed signatures of positive selection (Gloss et al., 2014). Our results therefore suggest that comparable phenomena have been acting at the level of polyphagan beetles. The CEs also show a statistically significant enrichment for positive results compared to the background, further supporting the diet detoxification hypothesis. The other positive results include a P450 OG and a CYS OG, neither of which led to a category enrichment compared to the background. The P450 OG is by far the largest among the positive results (Supplementary Figure 3), highlighting the importance of P450s in beetle (and generally insect) physiology with diverse roles beyond detoxification, e.g. hormone biosynthesis (Kong et al., 2014). However, while considered as significantly expanded and under selection by our model, the actual mean values in the suborders are not dramatically different. Importantly, the enrichment of positive results among candidates still holds if this OG is excluded (see Supplementary Results). The involvement of P450s in many other processes may explain why a broader difference between the suborders was not identified. Apart from one positive result among the cysteine proteases (no significant category enrichment), our study did not highlight additional expansions in other digestive enzymes or in transporters within a suborder. The lack of evidence for expansion in Polyphaga with respect to ABC transporters, which is the candidate functional category encompassing the highest number of OGs, may indicate that the ancestral diversity of transporters was sufficient for maintaining the excretion of toxins, despite variations in the substrates imposing a selective pressure on early stages of the detoxification pathway. Alternatively, if such pressure were acting on later stages of the pathway, i.e. transporters, its strength could have been too low for the detection power of our methods and data, unlike for GSTs or CEs.

## Conclusions

By comparing the degree of expansion among gene families involved in detoxification of plant secondary compounds in two suborders of beetles characterized by generally contrasting trophic niches (i.e., Polyphaga contain a high proportion of phytophagous species while Adephaga encompass mostly predacious species), we provide molecular genetic evidence supporting the popular hypothesis that Coleoptera species richness may be in part explained by their interaction with land plants. Candidate OGs of GSTs, CEs, P450s, and CYSs tested positive for adaptive LSEs in the phytophagous polyphagan beetle lineage, and categories of GSTs and CEs in particular, were enriched for OGs with such adaptive LSEs. Moreover, across all OGs tested, Polyphaga exhibited significantly more adaptive LSEs than Adephaga. This indicates that genes other than the candidate detoxification and digestion enzymes, which could include genes with functions less obviously related or unrelated to phytophagy, are also likely to have played a role in the adaptive success of Polyphaga. While this suggests that additional functional categories remain to be explored, contrasting gene family evolution across the two major suborders of beetles demonstrates a role for interactions with plant secondary compounds, and supports a role for phytophagy in general, as important drivers of the remarkable radiation of polyphagan beetles.

## Methods

A chart summarizing the main steps of the analysis is available as Supplementary Figure 10

### Data sources

This study included six genomes and 13 transcriptomes representing a balanced sampling of polyphagan and adephagan beetles, along with one representative of the sister group to Coleoptera, Strepsiptera, to root the species phylogeny. Annotated gene sets from four genomes were sourced from the i5k pilot project datasets (Robinson et al., 2011) (*Anoplophora glabripennis* v0.5.3 (McKenna et al., 2016), *Leptinotarsa decemlineata* v0.5.3 *(Schoville et al., 2018), Onthophagus taurus* v0.5.3*, Agrilus planipennis* v0.5.3) and two were independently published *Dendroctonus ponderosae* Ensembl Metazoa v1.0 (Keeling et al., 2013) and *Tribolium castaneum* Ensembl Metazoa v3.22 (Tribolium Genome Sequencing Consortium et al., 2008). One Polyphaga transcriptome, *Laparocerus tessellatus* (Supplementary Methods), was sequenced for this project and the others were provided by the 1KITE project (Supplementary Methods, http://www.1kite.org, Misof et al., 2014, Peters et al., 2017). A detailed list is presented in Table 1.

### Coding sequence predictions, transcriptome and genome quality assessments

Coding sequences and peptide sequences were predicted from all transcriptomes using TransDecoder (v2.0.1 https://transdecoder.github.io [last accessed May 4th, 2017]) along with a custom python script to retain the best-scoring entry among overlapping predictions. The coding sequences and peptide sequences from the genomes were retrieved from their official annotated gene sets. All genome and transcriptome gene sets were assessed using BUSCO (v2.0, python 3.4.1, dataset insecta_odb9/2016-10-21, mode proteins) (Waterhouse et al., 2018). This tool identifies near-universal single-copy orthologs by using hidden Markov model profiles from amino acid alignments. CD-HIT-EST v4.6.1 (Li and Godzik, 2006) was run on the protein sequences with a 97.5 percent identity threshold to ensure that all species datasets were filtered to select a single isoform per gene.

### Orthology delineation

The OrthoDB (Kriventseva et al., 2015) hierarchical orthology delineation procedure was employed to predict orthologous protein groups (OGs). Briefly, protein sequence alignments are assessed to identify all best reciprocal hits (BRHs) between genes from each pair of species, which are then clustered into OGs following a graph-based approach that starts with BRH triangulation. The annotated proteins from the genomes of *A. planipennis*, *O. taurus* and all transcriptomes were mapped to OrthoDB v8 at the Arthropoda level (with 87 species including four of the beetles with sequenced genomes). Mapping uses the same BRH-based clustering procedure but only allows genes from mapped species to join existing OGs. These OGs were then filtered to identify the 9,720 OGs with representatives from both Polyphaga and Adephaga to focus the study on OGs with evolutionary histories traceable to the last common ancestor of all the beetles, i.e. 5,188 OGs with genes from only one of the two suborders were removed.

### Species phylogeny

To build an ultrametric phylogeny required for the CAFE analyses, the maximum likelihood molecular species phylogeny was first estimated based on the concatenated superalignment of orthologous amino acid sequences from each of the datasets. Protein sequences of single-copy BUSCO genes and the best-scoring duplicated genes present in all species were individually aligned for each set of BUSCO-identified orthologs using MAFFT with the --auto parameter (Katoh and Standley, 2013) and each result was manually reviewed to exclude poor-quality alignments. Four hundred and five alignments were retained out of 436 and concatenated into a superalignment, partitioned according to the best model for each set of orthologs using aminosan 1.0.2015.01.23 (Tanabe, 2011). RAxML v8.1.2 (-f a -m PROTGAMMA -N 1000) (Stamatakis, 2014), and used to compute the maximum likelihood tree. The monophyly of Geadephaga and Hydradephaga was constrained to match the generally accepted resolution of Adephaga (as in McKenna et al., 2015). The chronos function of the R package ape (v3.4 on R 3.2.1, relaxed model) (Paradis et al., 2004) was used to obtain an ultrametric tree and the tip to root length was adjusted to match the approximately 250 million year evolutionary history of crown group Coleoptera (McKenna et al., 2015).

### Functional annotation and definition of candidate genes

InterProScan was run on all species protein sets (-appl Pfam --goterms, 5.16.55) (Jones et al., 2014) to identify protein families. Additionally, blastp 2.3.0 (Altschul et al., 1997; Camacho et al., 2009) was run against uniref50 (version Jun 22, 2016; Suzek et al., 2015) with an e-value cut-off of 1e-20. An OG was included in the set of candidate OGs when it had a match to both the uniref50 clusters and Pfam families (Finn et al., 2016) or gene ontologies (The Gene Ontology Consortium, 2017) as detailed in Table 2.

### CAFE analysis

The number of genes in OGs for each species were counted. All candidates and remaining (control) OGs were pooled together and processed with CAFE 3.1 (Han et al., 2013), to infer gene family evolution in terms of gene gains and losses. First, the python script provided by CAFE was used to estimate the error in our dataset. The CAFE software was then run using the mode in which the gain and loss rates are estimated together (λ) and a second mode in which they are estimated separately (gains=λ, losses=μ). The more complex model was retained as it reached a significantly better score (−199,989 for a single estimated parameter and −199,981 for two distinct estimated parameters, 2x delta log-likelihood = 16, chi-squared distribution, df=1). For the entire analysis, the CAFE overall p-value threshold was kept at its default value (0.01). To run CAFE on each suborder separately, the newick file was pruned to retain only required species using newick utils 1.1.0 (Junier and Zdobnov, 2010).

### Evolutionary models

To evaluate adaptive OG expansion, the likelihood of the count data was tested by optimizing parameters considering two methods provided by the OUwie R package v1.51 (Beaulieu and O’Meara, 2016; Beaulieu et al., 2012). First, a Brownian motion (BM) approach was used, which assumes no selection and thus differences result from a stochastic process whose rate is estimated. Second, Ornstein-Uhlenbeck (OU) models were used. They take into account an optimal family size which is obtained by selective pressure. Two groups were defined in the phylogeny, namely Polyphaga and Adephaga, to which the two different regimes to consider were assigned, plus a third regime to the root. This represents a simplified scenario allowing for the comparison of gene contents between one group and the other rather than attempting to estimate ‘levels’ of phytophagy or zoophagy across the phylogeny. The models BM1 (Brownian motion with a single rate for the whole tree), BMS (Brownian motion with different rates for each group), OU1 (selection towards the same optimum for both groups) were optimized as null hypotheses (H0) and compared to OUM (selection towards two optima, same variance) and OUMV (selection towards two optima, two variances) models as alternative hypotheses (H1). The Akaike information criterion corrected for sample-size (AICc) (Hurvich and Tsai, 1989) was used to compare models and an AICc>2 between the best H0 and the best H1 model was considered as significant to prefer the H1 model.

### Statistical enrichment

All results for candidates and controls were pooled together to obtain a background distribution of positive and negative results. Positive results are those OGs that passed the OUwie analysis, and negative results are all of the 9,720 OGs that did not obtain a significant overall CAFE p-value or did not pass the OUwie analysis. Then, 100,000 random draws (using the R function sample, without replacement) having the sample size of the candidate category to test for enrichment were taken from the background and the significant outcomes for Polyphaga and Adephaga were counted. A p-value was calculated for each group as follows: the number of random draws reaching the amount of significant outcomes found for the candidate category, or more, divided by 100,000. Additionally, the multiple tests conducted on each individual candidate category were corrected for false discovery rate (FDR) using the R p.adjust function (method BH, Benjamini Hochberg).

### Gene trees

The alignments for the gene trees were produced using MAFFT with the --auto parameter. The gene trees were computed with RAxML v8.1.2 (-f a -m PROTGAMMALGF -N 100) and plotted with EvolView (He et al., 2016; Zhang et al., 2012).

## List of abbreviations

ABC: adenosine triphosphate-binding cassette transporters
AICc: Akaike Information Criterion (small-sample-size corrected)
BM: Brownian motion
BUSCO: Benchmarking Universal Single-Copy Ortholog assessment tool
CAFE: Computational Analysis of gene Family Evolution tool
CE: carboxylesterase
CYS: cysteine protease
FDR: false discovery rate
GH: glycoside hydrolase
GST: glutathione S-transferase
LSE: lineage specific expansion
MAFFT: Multiple Alignment using Fast Fourier Transform tool
OG: orthologous group
OU: Ornstein-Uhlenbeck
P450: cytochrome P450 monooxygenase
SER: serine protease
UGT: uridine 5’-diphospho-glycosyltransferase

## Availability of data and material

The datasets generated and/or analyzed during the current study are available from the National Center for Biotechnology Information, https://www.ncbi.nlm.nih.gov, Genome Database (genomes and annotated genes) and Transcriptome Shotgun Assembly Sequence Database (transcriptomes). Details, including accessions, of all 1KITE transcriptomes are given in Supplementary Table 4 (currently unpublished transcriptomes will be released upon acceptance of this manuscript for publication).

## Funding

This work was supported by the United States National Science Foundation (DEB 1355169) and the United States Department of Agriculture (cooperative agreement 8130-0547-CA) to DDM, the Spanish grant CGL2013-42589-P, awarded by the MINECO and co-financed by FEDER to BCE, the Science Foundation DFG grant BA2152/11-1, 2, the BGI-Shenzhen, the China National Genebank, and the following Swiss National Science Foundation grants: 31003A_143936 (PI), 31003A_173048 (MRR), PP00P3_170664 (RMW), PP00P3_144870 (NA) and PP00P3_172899 (NA). Funding for open access charge: Museum of Natural History of Geneva.

## Author contributions

MS and NA conceived the study. PI delineated orthology. CP prepared the libraries for *Laparocerus tessellatus* transcriptome sequencing. HEE, DDM, BM, SS, and XZ provided access to 1KITE transcriptome data. MS conducted the analyses. BCE, MRR, JR, RMW and NA supervised the analyses. MS, RMW, and NA wrote the manuscript, with input from all authors.

## Acknowledgements

The authors thank Rolf Beutel, Adam Slipinski, Kai Schuette, Ralph Peters, and Eric Anton for providing specimens for the 1KITE samples, as well as Michael Balke, Jia Fenglong, Xu Shengquan, and Alexandros Vasilikopoulos for the *Sinaspidytes wrasei* data. The authors also thank Shanlin Liu, Alexander Donath, Lars Podsiadlowski, Karen Meusemann, and the 1KITE Coleoptera group for providing transcriptome assemblies of 1KITE species. The authors gratefully acknowledge pre-publication access to gene predictions for *Onthophagus taurus* and *Agrilus planipennis* from the Baylor College of Medicine Human Genome Sequencing Center’s i5k pilot project, represented by Stephen Richards. The authors additionally thank Catherine Berney who contributed to the production of the *Laparocerus tessellatus* transcriptome, and Nicolas Salamin for advice on the applied methods and feedback on the manuscript. Some of the computations were performed at the Vital-IT (http://www.vital-it.ch) center for high-performance computing of the Swiss Institute of Bioinformatics.

